# Lateral Cerebellar Nucleus Lesions Disrupt Rule-Based Category Learning and Cognitive Flexibility in Rats

**DOI:** 10.64898/2026.02.11.705339

**Authors:** Shannon L. Wachter, Matthew B. Broschard, Krystal L. Parker, John H. Freeman

**Author notes:** **Correspondence:** John H. Freeman Krystal L. Parker.

## Abstract

Cerebellar communication with the prefrontal cortex (PFC) may play a significant role in cognitive functions. Our previous studies found that rule-based (RB) category learning depends on the PFC in humans and rats. The PFC is also crucial for behavioral flexibility following rule-changes in various tasks. Very little is known regarding the role of the cerebellum in RB category learning. The current study was designed to determine whether the cerebellum plays a role in RB category learning, and in categorization following a rule switch. Female and male rats were given bilateral lesions of the lateral cerebellar nuclei (LCN) or a control surgery and trained on an RB category learning task followed by a category rule switch. A subset of rats was trained on a control discrimination task with the same trial procedures as the categorization task. Rats with LCN lesions took significantly longer to learn both the first and second category rules but were not impaired on the control task. Computational modeling revealed less task engagement and increased switching between engaged and non-engaged states in the LCN lesion group. Several measures of task performance indicated that the category learning deficit was not caused by a motor impairment, response bias, or an inability to discriminate the stimuli. The category learning deficits with LCN lesions were related to reduced accuracy of stimulus classification, an inability to maintain task engagement, and loss of flexibility. The results show, for the first time, that the cerebellum plays a crucial role in category learning and category rule-switching.

---

The cerebellum was traditionally thought to be primarily involved in motor planning and execution. However, it is now widely acknowledged that it also contributes to cognitive processes [1, 2, 3]. Neuroimaging studies have shown changes in cerebellar activity during cognitive tasks [1, 4–6]. Patients with cerebellar damage have deficits in cognitive functions such as working memory, selective attention, and rule-switching [7–9]. Cognitive deficits in schizophrenia and other neuropsychiatric disorders are associated with altered cerebellar activity [10, 11]. The cognitive deficits associated with cerebellar lesions and neuropsychiatric disorders have been attributed to disrupted communication between the prefrontal cortex (PFC) and cerebellum [10, 11, 12].

The cerebellar nuclei receive distinct input from Purkinje cells in the cerebellar cortex (e.g., Crus I to the lateral cerebellar nuclei (LCN)) [13]. Each nucleus has different pathways to forebrain structures, and thereby contribute to different functions [13, 14, 15]. The LCN projects to the medial PFC via the thalamus and these connections are thought to contribute to executive functions [16–19]. This hypothesis is supported by optogenetic and pharmacological studies showing that disruption of cerebellar output from the LCN or changes in the activity of Purkinje cells that project to the LCN have deleterious effects on performance in interval timing and spatial working memory in rodents [20, 21, 22]. Thus, there is growing evidence that the cerebellum mediates cognitive functions through di-synaptic projections from the LCN to the PFC.

Categorization is a task that requires organisms to form equivalence classes based on perceptual or functional similarity. Categorization involves multiple executive functions such as selective attention, memory, and decision-making for humans and rats [23–29]. Previous studies found that the PFC is essential for rule-based (RB) category learning in which subjects selectively attend to a single perceptual dimension or rule (e.g., spatial frequency, orientation, color, size, brightness) among multiple dimensions to categorize stimulus exemplars. The PFC is not necessary for information integration (II) category learning, where subjects must use both dimensions to properly categorize [24]. Rats and humans with PFC damage have difficulty consistently using the relevant or optimal rule to guide decisions regarding category membership of exemplars in RB learning [24,29]. Because of the LCN-to-PFC connectivity, we predict that lesioning the LCN will also impair RB category learning by disrupting connectivity to the PFC.

Rule switching in an RB categorization task can be used to assess cognitive flexibility. RB categorization tasks utilize selective attention to a single stimulus dimension and therefore, switching the rule to categorize involves shifting attention from one dimension to the other [23–28]. When the former optimal rule becomes the suboptimal rule, subjects experience interference similar to interference seen with the Stroop Task [30]. Since the Stroop Task and other rule-switching tasks require the PFC, we can make inferences regarding the status of PFC-dependent cognitive flexibility by switching the RB rule after initial category learning. Thus, disrupting LCN-to-PFC communication might impair performance following a rule switch in RB categorization.

Two prior studies attempted to address the role that the cerebellum plays in RB and II categorization in patients with cerebellar damage; however, they did not find a significant difference from controls [31, 32]. These studies used patients with a large range of lesion sizes and locations; very few had lesions in the posterior lateral hemispheres, and even fewer had bilateral lesions. Therefore, it is important to examine the role of the cerebellum in category learning using an animal model where damage can be more precisely localized.

In this study, we examined the effects of LCN lesions on RB category learning and flexibility following a category rule switch in rats. We utilized an RB categorization paradigm where rats were presented with a stimulus containing black and white gratings that vary in spatial frequency and orientation (Fig. 1). The rats need to selectively attend to a relevant dimension (spatial frequency or orientation) and ignore the irrelevant dimension until they reached an accuracy criterion [23–28]. The relevant dimension was then switched to the formerly irrelevant dimension to test cognitive flexibility. We hypothesized that lesioning the LCN would disrupt performance in RB categorization and impair cognitive flexibility by severing cerebellar communication with the PFC.

**Figure 1.**
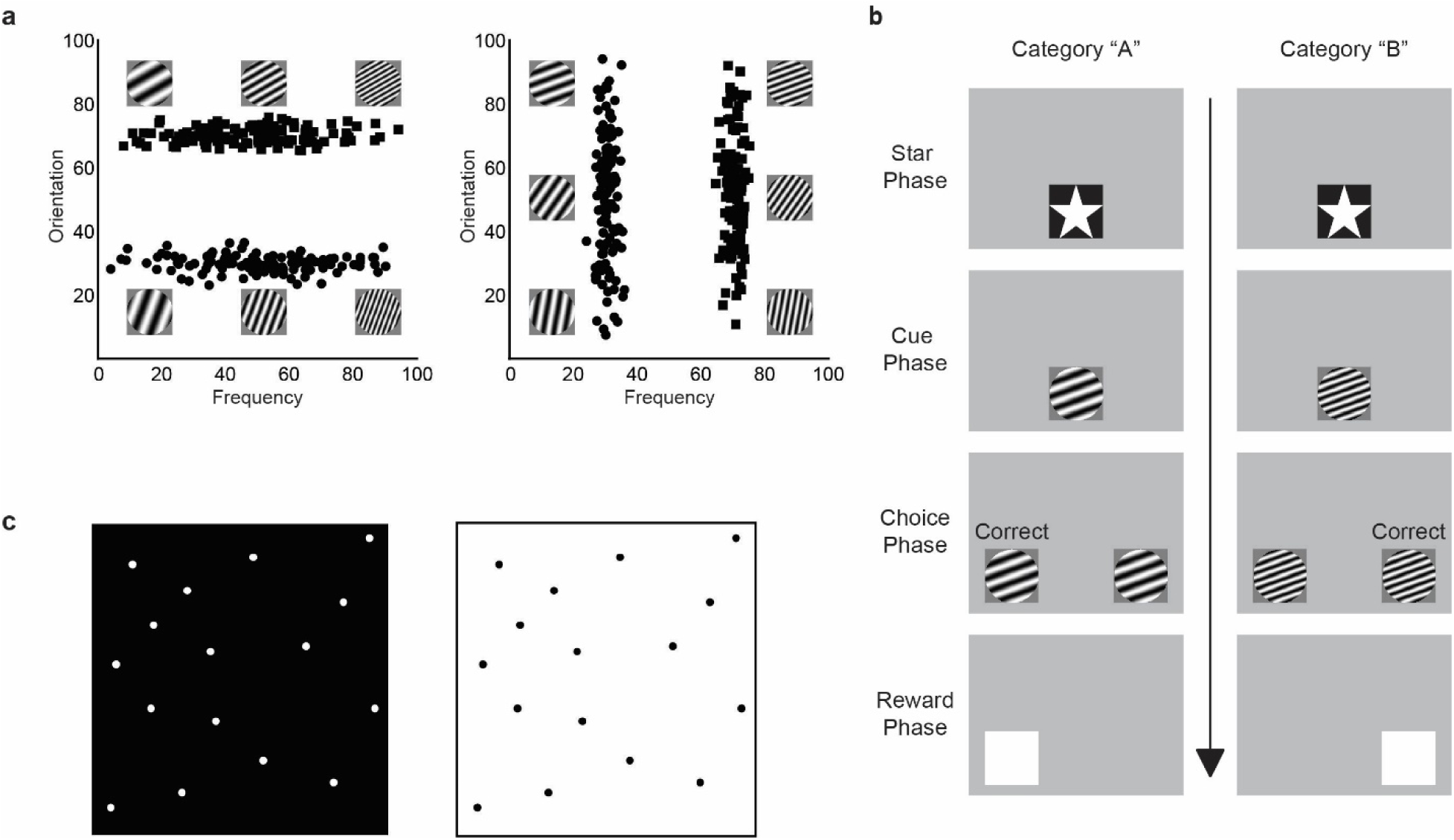
Category learning and control tasks. a, Rule-based category learning task stimulus distributions and example stimuli for the orientation rule (left) and spatial frequency rule (right). b, Trial sequence and procedures for the rule-based category learning task. First, rats touched a star to start the trial, then an exemplar is presented in the middle of the screen (Cue phase). After touching the exemplar three times, copies of the exemplar appeared on the left and right sides as response keys (Choice phase). Rats touched either response key, if the category selection was correct then a white box appeared (Reward phase). The white box served as a secondary reinforcer. c, Control task stimuli. Rats were trained to discriminate a white box from a black box using the same procedures as the categorization task except that there was only one stimulus in each category. The discrimination task controls for all aspects of the task other than classifying stimuli from distributions.

## Materials and methods

### Subjects

Thirty-two Long Evans rats (14 females and 18 males) were used. Rats were individually housed and kept on a 12-hour light-dark cycle with training occurring during the light cycle. Rats were given water *ad libitum*, but food was restricted throughout the experiment. Weights were recorded daily to ensure rats maintained no less than 85% of their free-feeding weight (250–350g). Procedures were approved by the Institutional Animal Care and Use Committee at the University of Iowa. See [23–28] for more methodological details.

### Surgery

Rats were anesthetized with isoflurane (5%), then placed in the stereotaxic apparatus, where isoflurane was maintained between 1% to 4% throughout the duration of the surgery. After the skull was exposed, Burr holes were drilled through the skull at AP: −11.4, ML: ± 3.6, DV: −5.7 for males and AP: −11.1, ML: ± 3.2, DV: −5.5 for females. Bilateral electrolytic lesions were made by delivering DC current (1.0 mA for 15 seconds) through a stainless-steel insect pin (size 00, insulated with Epoxylite) via a constant current lesion maker (Ego Basile 3500). Rats assigned to the sham-operated control group underwent the same surgical procedure but without electrical stimulation. Meloxicam (1.0 mg/ml) was administered during surgery and within 24 hours after surgery for analgesia. Rats recovered for a week before beginning pre-training procedures.

### Touchscreen Apparatus

For all shaping and experimental sessions, rats were placed within custom-built touch-screen chambers (36 x 41 x 36cm). The chambers contained an LCD flat-screen computer monitor (Model 1550V, NEC, Melville, NY) mounted on one wall which presented visual stimuli to the rats. Rats interacted with the monitor through a touchscreen (15in, EloTouch Systems Fremont, CA) placed in front of the monitor. The wall opposite the monitor, holds a food tray (6.5 x 13 x 4.5cm) where 45mg grain pellets (LabTab MLab Rodent, TestDiet, IN) were delivered via a rotary pellet dispenser (Med Associates Inc., Georgia, CT, model ENV-203IR) which was controlled through an electrical board (Model RS-232, National Control Devices, Osceola, MO). Above the food tray is a house light which was turned on at the onset of a trial during shaping and experimental sessions. Outside the operant chamber, but within the behavioral room, white noise played during sessions to minimize distractions. All experimental tasks and procedures were controlled through a custom MATLAB script (MathWorks, Natick, MA).

### Pre-training procedures

After recovery from surgery, rats were handled daily for three days. Once handling was completed, each rat underwent a pseudo-open field task using the surface of a standard laboratory cart to further habituate them to the experimenter and environment. During the open field procedure, rats were presented with twenty 45mg grain pellets scattered across the surface of the cart and given 20 min to consume all twenty grain pellets. This procedure continued daily for each rat until they consumed all grain pellets within the allotted time. Once completed, rats were shaped daily to interact with a touchscreen within an operant chamber. Shaping included three separate phases, with each phase progressing towards a similar trial sequence as seen in category training [23–28]. By the end of shaping, the rats were using the same trial procedure as the category training sessions, but without the stimuli used during categorization.

### Category stimuli and category rules

Category stimuli (239 x 239 pixels) contained black and white gratings that varied along two continuous dimensions, spatial frequency and orientation. Spatial frequency of gratings ranged between 0.2532 cycles per visual degree (cpd) to 1.2232 cpd. Orientation of the gratings ranged from 0 rad to 1.75 rad. These dimensions were transformed to fit within a common range (i.e., 0 to 100; see [23–25] for more details). The given ranges of spatial frequency and orientation are within perceptual limits of rats [33].

Two categories were created by placing two bivariate normal distributions within the normalized space (Fig. 1a, Category A: Xμ = 30, Yμ = 50, Xσ = 2.5, Yσ = 20; Category B: Xμ = 70, Yμ = 50, Xσ = 2.5. Yσ = 20) where each point within that normal distribution is a representative category stimulus. Rats learned RB categorization where each rule is determined by category distributions perpendicular to one axis (i.e., Fig. 1a) where the other rule was created by turning the distributions 90 degrees [25].

### Category training and rule switching

Each rat received training on both frequency and orientation rules. Starting rules were randomly assigned to each rat. Rats were trained to a criterion of 75% correct for both category A and B for two consecutive training sessions or until that reached a maximum of 30 sessions. Each training session contained 80 training trials (Fig. 1b). On each trial, a star stimulus was presented in the center of the screen (trial start). After the rat touched the star, a category stimulus, chosen from the training distribution, replaced that star stimulus (cue phase). After the rat touched the category stimulus three times, that same category report keys were then presented on the left and right sides of the screen (choice phase). Categories are mapped spatially, where the left side represented category ‘A’ while the right side represented category ‘B’. Rats selected either the left or right stimulus to indicate which category they thought that category stimulus belongs to. If the correct side was chosen, a white box appeared on that side which when pressed delivered a food reward (reward phase). If the incorrect side was chosen, the trial would repeat with the same category stimulus. After a 5 to 10 second timeout, the trial was repeated from the cue phase. This cycle repeated until the correct response was chosen.

Once a rat had reached a criterion of 75% for both categories for two consecutive training sessions or had 30 training sessions, the category relevant dimension (rule) was switched (e.g., orientation to frequency). Now the once relevant dimension has become the irrelevant dimension. Training continued using the same trial format as described above, with new distributions perpendicular to the previous ones. The criterion and maximum number of sessions remained the same for the second rule.

### Simple discrimination

To ensure that the differences between groups were specific to categorization, rats were given training sessions to learn a simple discrimination task after category training. Rats were trained to differentiate between two images - a light box and a dark box, both with a spread of dots to increase stimulus complexity (Fig. 1c; Kim, Castro, Wasserman, & Freeman, 2018). Trial procedures were similar to category training, except instead of choosing category representation, the stimuli were mapped to either the left or right report key. Each session contained 72 trials, and training continued until the rats reached 75% accuracy for both images for two consecutive sessions.

### Histology

After behavioral testing, rats were perfused and lesion placement was confirmed. Rats were given a lethal dose of sodium pentobarbital (Fatal-Plus) and then perfused with PBS followed by 10% formalin. Brains were removed and stored at 4□ in a solution of PBS and 30% sucrose. Using a cryostat, 50μm coronal slices were collected through the entire deep cerebellar nuclei (DCN). Slices were then plated and stained with thionin. Pictures of the tissue were collected using a light microscope. Lesion size was quantified using ImageJ. Lesion volume was calculated using the equation of a sphere with the radius of the sphere taken from the section containing the largest lesion. Rats with lesions significantly outside the LCN were excluded from analysis.

### Statistical analysis

Non-parametric methods were used to test within-subject and between-subject differences (Wilcoxon signed rank test and Mann-Whitney U Test, respectively) on sessions to criterion (R, version 4.4.0). A non-parametric test was used due to the non-normal nature of the lesioned data, where several rats in Rule 2 reached our criterion cut-off of 30 sessions. Other dependent measures (accuracy and reaction time) were analyzed using linear mixed effects modeling. Models for training sessions included fixed effects for experimental group, rule, training session, and a quadratic function across training sessions and random effects for intercept, slope and the quadratic function. Sex differences were assessed by including a sex covariate in the models. A model simplification strategy was used to find the maximal random effect structure justified by the data [34].

### Iterative Decision Boundary Modeling

Behavioral strategies were characterized using iterative decision boundary modeling [35, 36]. Through this method, multiple models are iteratively tested across trials to determine how different category responses (i.e., category A vs B) are mapped onto the stimulus space. Fitting these models to the rats’ behavior allowed for inferences to be made about which strategy was used to make a category decision and when the rats switched strategies. For this analysis, two boundary types were used, including two unidimensional models and a bidimensional model. Unidimensional models created boundaries using a single stimulus dimension (i.e., spatial frequency or orientation). Unidimensional models could be characterized as either relevant (the boundary following the relevant rule, which is optimal) or irrelevant (the boundary following the irrelevant rule, which is suboptimal). The bidimensional model created boundaries by combining information from both stimulus dimensions, which is considered a suboptimal strategy throughout this task. These models were compared to a control model, which did not create a decision boundary, but instead assumed the rats were randomly guessing. See [36] for more details.

For the analysis in the following experiments, four boundary types were used, including two unidimensional models, a bidimensional model, and a control model. Unidimensional models created boundaries using a single stimulus dimension (i.e., spatial frequency or orientation). Unidimensional models could be characterized as either relevant (the boundary follows the relevant rule, which is optimal) or irrelevant (the boundary follows the irrelevant rule, which is suboptimal). The bidimensional model created boundaries by combining information from both stimulus dimensions, which is considered a suboptimal strategy throughout this task. These models were compared to a control model, which did not create a decision boundary suggesting the rats were randomly guessing.

This computational modeling not only determines which strategy is being used but also can detect the switches between strategies [35, 36]. In this approach, a set of “switch models” are fit to the data in addition to the four models described above, referred to as “basic models”. The switch models assume that within a particular window of data, the best-fitting model switches from one basic model to another, creating 12 possible switch strategies. In total, sixteen models are fit to each window.

For each rat, the full set of models were fit to the first 100 training trials using the MATLAB function *fmincon*. Bayesian information criterion (BIC) values were calculated and compared for each model where the model with the smallest BIC value was determined to be the best fit (Hélie et al., 2017). Subsequent fittings increased the number of trials included by one trial until the algorithm detected a switch in strategy. If the same switch model was the best-fitting model for the last 15 iterations and the estimated switch was consistent within a range of 21 trials for the last 15 iterations, then the switch in strategy was recorded. The trial in which this switch occurred was derived from an average estimated switch trail from the last 15 iterations. Once this was determined, the iterations resumed. Iterations continued until it reached the end of the training trails. A Chi-Square test as used to determine differences between groups.

### Engaged and Non-Engaged Trials

Decision boundary modeling was also used to determine the rat’s engagement in the task through detecting local changes in strategies [35, 36]. The transition from engaged to non-engaged trial sections was determined from the best-fitting model switching from a boundary model (uni- or bidimensional) indicating engagement to a control model indicting guessing, and back to engagement (i.e., a uni- or bidimensional model). Models were fitted to small sections of the rats’ choice data (i.e., 20 trials per iteration; the fitting window shifted by 1 trial between iterations). For each iteration, we calculated the Likelihood Ratio Test [38] between the model fits. Specifically,

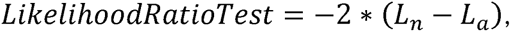

where *L_n_* is the log-likelihood of the null model (i.e., the RGM) and *L_a_* is the log-likelihood of the alternative model (i.e., the optimal decision boundary model). This statistic follows the chi square distribution, where the degrees of freedom are equal to the number of additional parameters in the alternative model compared to the null model (in this case, 1). Thus, for α = 0.05, any value above 3.841 was considered statistically significant. Iterations continued until the end of training was completed. The ‘Engaged’ trial sections were defined as consecutive iterations in which the Likelihood Ratio Test was significant. To minimize type I errors, only trial sections larger than 15 iterations were recorded.

## Results

### Histological assessment of LCN lesions

Using a light microscope, each lesion was examined and imaged. LCN boundaries were determined with a stereotaxic atlas [39]. On average, bilateral lesions extended between −10.65 mm and −11.78 mm AP (Fig. 2), with the average electrode placement at −11.22 mm from bregma. The mean lesion area was 1.681mm^2^ on the right and 1.789mm^2^ on the left. Lesions in 4 male rats, extended farther posteriorly than the others and lesions of 1 female extended farther anteriorly. Cerebellar cortical damage only occurred when lesions extended either farther posteriorly or anteriorly, with the average cortical lesion area being .744mm^2^ on the right and .819mm^2^ on the left. Cerebellar cortical areas that were partially lesioned included the lateral paramedian lobule and copula of the pyramis in rats with farther posterior lesions. The rat with anterior lesions had damage to the middle cerebellar peduncle. However, no behavioral differences were observed between rats with posterior or anterior damage when compared to those with lesions contained in the LCN boundaries. There was very little evidence of damage to the interpositus nucleus, with the very lateral edge of the interposed nuclei damaged in 3 rats.

**Figure 2.**
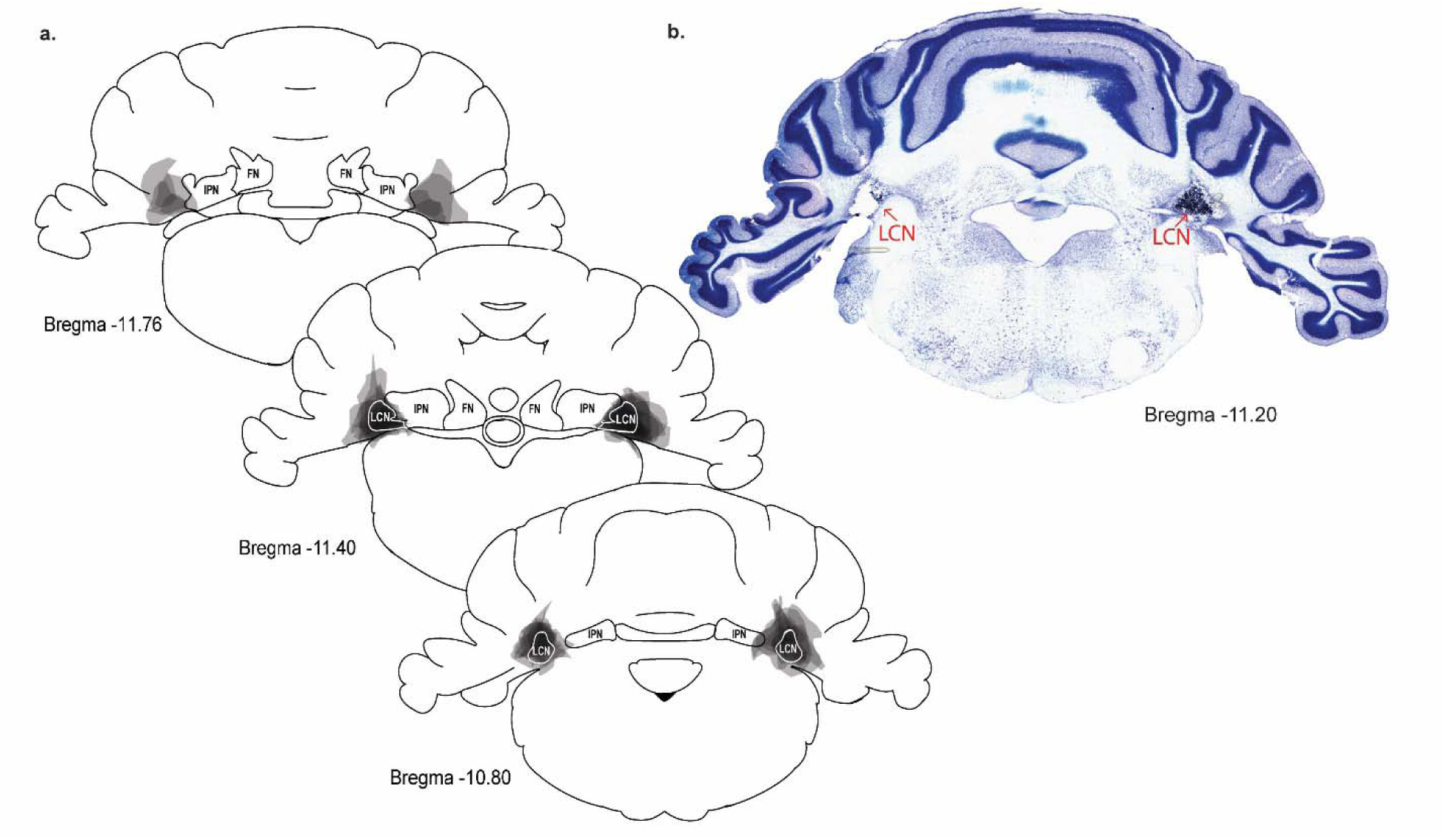
Lesion location and spread. a, Bilateral lesions are depicted with shaded areas for each rat to show the extent of overlap and spread. The lesions extended between −10.65 mm and - 11.78 mm AP, with the average electrode placement at −11.22 mm from bregma. The mean lesion area was 1.681mm^2^ on the right and 1.789mm^2^ on the left. b, Example thionin stained section showing bilateral damage to the LCN.

### LCN lesions impaired rule-based category learning and rule switching

Rats with bilateral LCN lesions learned the rule-based categorization task more slowly than the rats that underwent control surgery when the categories were based on spatial frequency or orientation (Rule 1, Fig. 3a). When the rule was switched (Rule 2), the lesion group was even more impaired (Fig. 3a). Session to criterion was assessed for normality using a Shapiro-Wilk test and a Levene’s Test to assess homoscedasticity. Sessions to criterion for the lesion group during Rule 2 was not normally distributed *(W* = .813, *p* < 0.01); however, variances of the groups were not significantly different from each other for both rules. Due to the non-normally distributed data, non-parametric tests were used, and medians were used to summarize sessions to criterion. Group differences in the median sessions to criterion were analyzed using Mann-Whitney U tests. The lesion group took significantly longer to reach criterion for Rule 1 (*U* = - 3.596, *p* < 0.001), and Rule 2 (*U* = −4.404, *p* < 0.001), compared to controls. The number of sessions to criterion was then compared for Rule 1 vs. Rule 2. All rats took longer to reach criterion when the rule was switched (*Mdn* = 17) than for the first rule (*Mdn* = 15). A Wilcoxon Signed-Rank test indicated that this difference was statistically significant (*z* = −2.023, *p* < 0.05). In addition, lesioned rats took longer to reach criterion in Rule 2 (*Mdn* = 21) than Rule 1 (*Mdn* = 20), *z* = 7.5, *p* < 0.05, while there was no difference between Rule 1 and Rule 2 for controls. These results indicate that LCN lesions impaired rule-based category learning and switching to an orthogonal rule (e.g., spatial frequency to orientation). The impairment in learning Rule 2 in lesioned rats suggests that the cerebellum may be necessary for the cognitive flexibility needed to switch from one rule to another.

**Figure 3.**
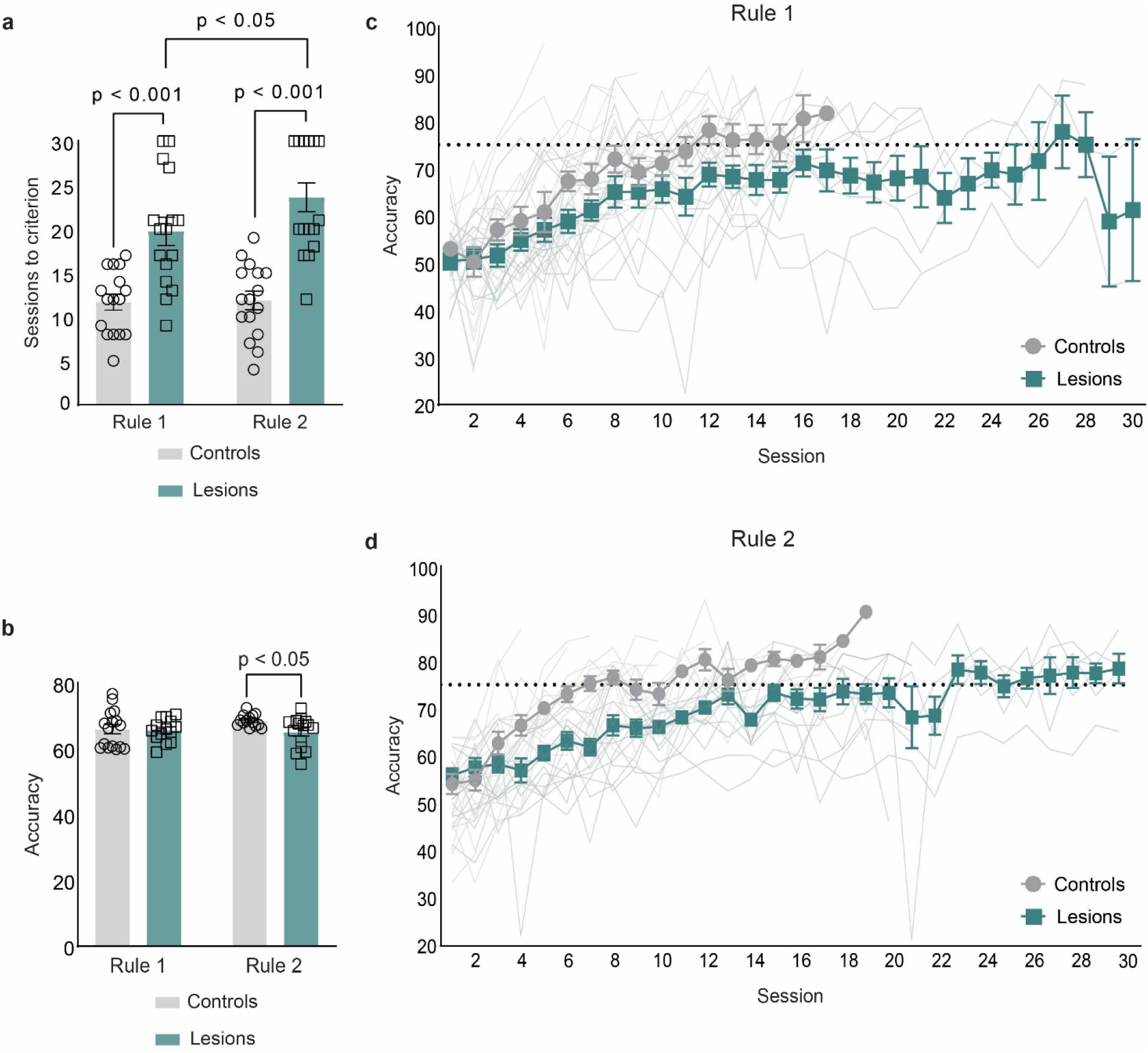
Category learning data for Rule 1 and Rule 2. a, Mean (± SEM) sessions to reach criterion (75% accuracy on both categories for two consecutive sessions) in the control group and group given LCN lesions. b, Mean (± SEM) accuracy across all sessions. c, Mean (± SEM) and individual (thinner lines) learning curves for Rule 1. d, Mean (± SEM) and individual learning curves for Rule 2. Horizontal dotted lines in c and d indicate the learning criterion.

There was no difference between controls and lesions in overall accuracy (Fig. 3b) throughout training for Rule 1 or Rule 2. A Mann-Whitney U test was used to assess group differences in overall median accuracy. There was no significant difference between rats with lesions and control rats for Rule 1 (*U* = 111 *p* = .983), but there was a significant difference for Rule 2 (*U* = 57, *p* < 0.05). A Wilcoxon Signed-Rank test was used to identify differences across rules within groups and was not statistically significant (*z* = 154, *p* = 0.413).

Learning curves were analyzed using mixed effects modeling. The full models included fixed effects for experimental group, training session, rule, a quadratic function across sessions, and random effect for intercept, slope, and the quadratic function. Accuracy increased significantly across sessions for all rats, *t*(675.34) = 6.887, *p* < 0.001. The rate of learning for Rule 2 was slower than in Rule 1 for all rats, *t*(809.9) = −2.034, *p* < 0.05. The rate of learning, as determined by accuracy, for lesioned rats was significantly slower than controls in Rule 2 compared to Rule 1, *t*(365.12) = 2.313, *p* < 0.05. When rule accuracy was analyzed separately, lesioned rats showed significantly lower accuracy compared to controls across sessions in Rule 2, *t*(454.77) = 4.30, *p* < 0.001, but not Rule 1, *t*(433.73) = −0.602, *p* = 0.547. These results indicate that rats with LCN lesions had difficulty learning overall, as determined by accuracy, but significance is driven by the lesioned rats’ inability to learn Rule 2 (Fig. 3d). According to linear modeling, lesioned rats were not different from controls in accuracy over sessions in Rule 1; however, there is a significant difference in how many sessions it took lesioned rats to reach criterion. Suggesting that although lesioned rats could get close to criterion at the same rate as controls, they were unable to maintain that accuracy long enough to successfully reach criterion (Fig. 3c).

### LCN lesions did not impair reaction time

Reaction time data during the choice phase can indicate slower decision-making or a motor deficit [24]. We therefore analyzed reaction times during the choice phase using a mixed 2×2 ANOVA (Fig. 4a). There was no group effect between controls and lesioned rats, *F*(1, 26) = 0.152, *p* = 0.700. However, there was a significant difference in choice reaction time between Rule 1 and Rule 2, *F*(1, 26) = 29.968, *p* < 0.001, _η_*²* = 0.156. The two-way interaction between group and rule during the choice phase was not significant, *F*(1, 26) = 0.00024, *p* = 0.988. Rats had slower reaction times while trained with Rule 1 (_μ_ = 1.33, _σ_ = 0.432) compared to training with Rule 2 (_μ_ = 0.98, _σ_ = 0.364). These results indicate that the rats with LCN lesions were not impaired in the time it took to make a decision and did not have a motor deficit.

**Figure 4.**
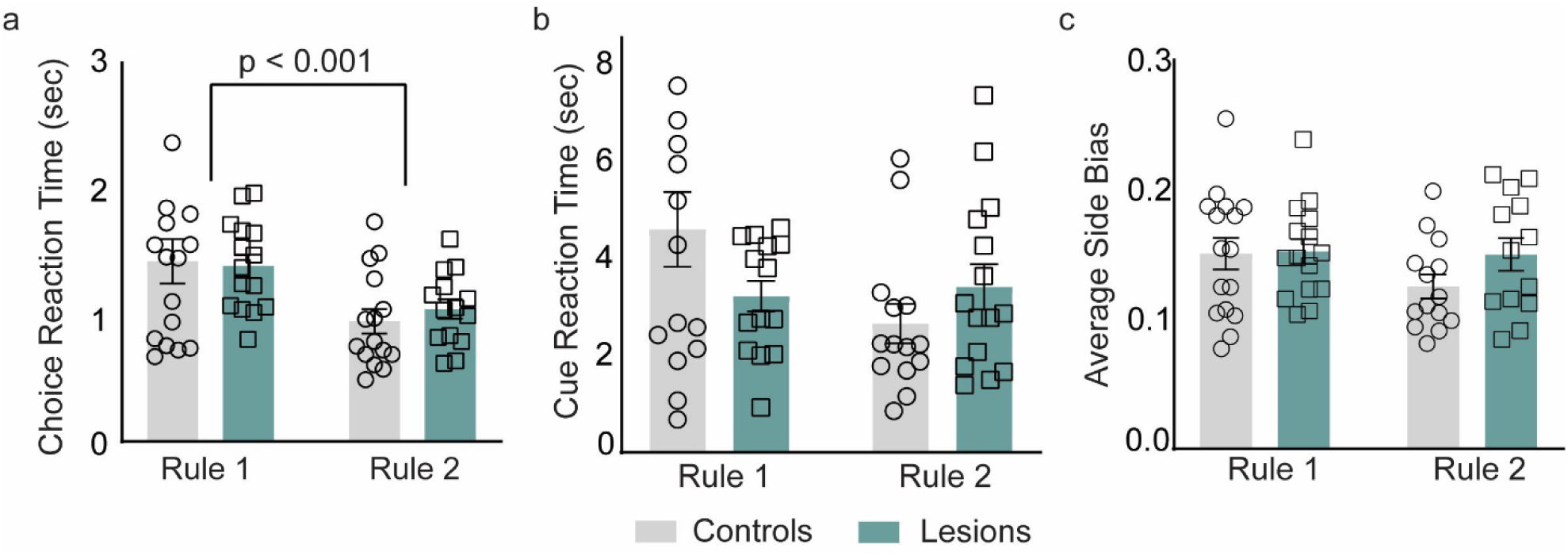
Choice reaction time, cue reaction time, and side bias data. a, Mean (± SEM) reaction time during the choice phase in the control group and LCN lesion group for Rule 1 and Rule 2. b, Mean (± SEM) reaction time during the cue phase. c, Mean (± SEM) side bias.

Linear mixed-effects modeling was used to analyze the choice reaction times across sessions to determine if there were effects during learning. When analyzed across sessions, no significant interactions between groups in reaction times were found during the choice phase (*t*(592.4) = −1.064, *p* = 0.288), indicating that the performance deficit during categorization could not be explained by a motor deficit or slower decision-making during the choice phase. The only significant finding was that choice reaction time decreased throughout training, *t*(551) = −3.87, *p* < 0.001.

Slower reaction times during the cue phase can indicate motor deficits or difficulty classifying the stimulus. Reaction time data during the cue phase were also analyzed using a mixed 2×2 ANOVA. (Fig. 4b). Similar to the choice phase, the two-way interaction between group and rule during the cue phase was not significant, *F*(1, 19) = 0.074, *p* = 0.788. In addition, the main effects of group (*F*(1, 19) = 4.199, *p* = 0.055) and rule (*F*(1, 19) = 1.377, *p* = 0.255) were not significant. The results of the reaction time data for the cue phase further indicate that rats with LCN lesions did not have a deficit in motor function or difficulty classifying the cue.

Linear mixed-effects modeling was used to analyze reaction time during the cue phase across sessions. There was no significant difference between lesioned and control rats in reaction time across sessions, *t*(745.37) = 2.4798, *p* = 0.12. Similarly, there was no significant difference between groups for either rule, *t*(734.15) = 0.31, *p* = 0.58. All other interactions were not significant for reaction times during the cue phase, indicating that the deficit in accuracy could not be explained by a motor deficit or difficulty classifying the cue.

### LCN lesions did not cause a response bias

A deficit in accuracy in a two-choice categorization task could be caused by a response bias in which that rats consistently choose one side during the choice phase (i.e., left or right). Response bias is the absolute value of the difference between left and right responses over the total number of responses [23]. Response bias was also analyzed using a mixed 2×2 ANOVA (Fig. 4c). No effects were found for the group x rule interaction, *F*(1, 28) = 3.88, *p* = 0.061; main effect of rule, *F*(1, 28) = 0.733, *p* = 0.399; or main effect for group, *F*(1, 28) = 4.096, *p* = 0.053. These results indicate that the deficit in accuracy in the LCN lesion group was not caused by a response bias.

Linear mixed effects modeling was used to analyze the response bias data across sessions. There was no significant difference in response bias between groups across rules (*t*(941.45) = −0.134, *p* = 0.893) further indicating that the deficit in accuracy was not caused by a response bias.

### LCN lesions do not impair strategy use and strategy switching

Humans switch strategies in categorization learning tasks until they find the optimal approach for the task at hand, but this ability is impaired in patients with damage to the prefrontal cortex [29]. Rats also switch strategies to the optimal strategy during category learning. Failure to switch strategies could account for the deficit in accuracy following LCN lesions. For visualization of these results, individual data were separated into 50 equal bins. Each bin represents the proportion of rats using each strategy (Fig. 5). The majority of controls (Fig. 5a) and lesioned (Fig. 5b) rats switched to the optimal unidimensional strategy while learning the first category rule (13 of 16 rats and 11 of 16, respectively). However, the optimal unidimensional strategy was not maintained through the end of training Rule 1, where rats with LCN lesions showed more random guessing (1 of 16) and bidimensional strategy use (7 of 16) than the controls (0 of 16 and 4 of 16, respectively) by the end of training with Rule 1. For Rule 2, very few controls (3 of 16 rats) or lesioned rats (4 of 16 rats) choose the optimal unidimensional strategy; instead, controls used a bidimensional strategy (12 of 16 rats), while lesioned rats were spread across random guessing (1 of 16 rats), irrelevant unidimensional (2 of 16 rats) and bidimensional (9 of 16 rats) strategies. The finding that controls relied on a bidimensional strategy during Rule 2 rather than the optimal unidimensional strategy suggests some retention of the previously learned rule while learning the second rule. Overall, there was no significant difference between groups in number of rats choosing the optimal strategy (*X^2^*(.4654, 1) = 0.682, *p* = 0.495) nor choosing the bidimensional strategy (*X^2^*(1.998, 1) = 1.414, *p* = 0.158). These results indicate that LCN lesions did not significantly affect strategy development during category learning or after the rule change.

**Figure 5.**
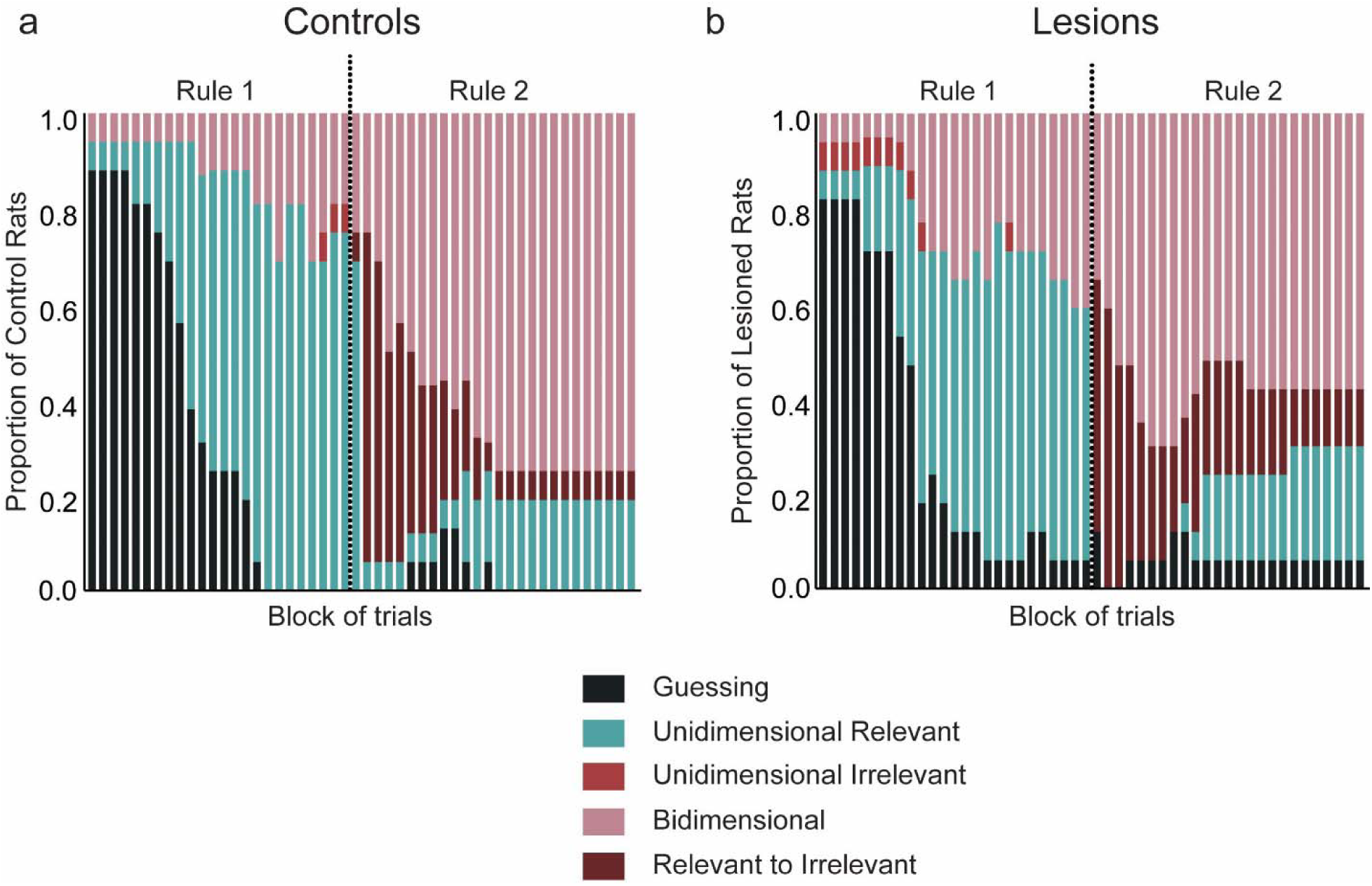
Iterative decision boundary modeling data. Proportion of rats in the control group (a, left) and LCN lesion group (b, right) exhibiting different strategies across blocks of trials. Strategies were classified as guessing (no model fit the data), unidimensional relevant (the correct category rule), unidimensional irrelevant (incorrect rule), bidimensional (using portions of both stimulus distribution), and relevant to irrelevant (maintaining the Rule 1 correct strategy into Rule 2 training). There were trends toward more incorrect strategies in the LCN lesion group, but there were not statistically significant differences.

### LCN lesions caused a decrease in task engagement and an increase in transitions between task engagement and non-engagement

Rodents and primates show periods of task engagement and non-engagement in various tasks [28, 36, 40–42]. Task performance could suffer from non-engagement and might account for the accuracy deficit in rats with LCN lesions. To address this factor, we analyzed the proportion of engaged and non-engaged trials (Fig. 6a), as well as switches between them with mixed ANOVAs. To compare the proportion of engaged trials across groups and rules, a 2×2 mixed ANOVA was used. No effects were found for the group x rule interaction, *F*(1, 59) = 0.8129, *p* = 0.371 and main effect of rule, *F*(1, 59) = 1.295, *p* = 0.26. However, there was a main effect for group, *F*(1, 59) = 9.343, *p* < 0.01, which was due to a reduction in engaged trials for the LCN lesion group.

**Figure 6.**
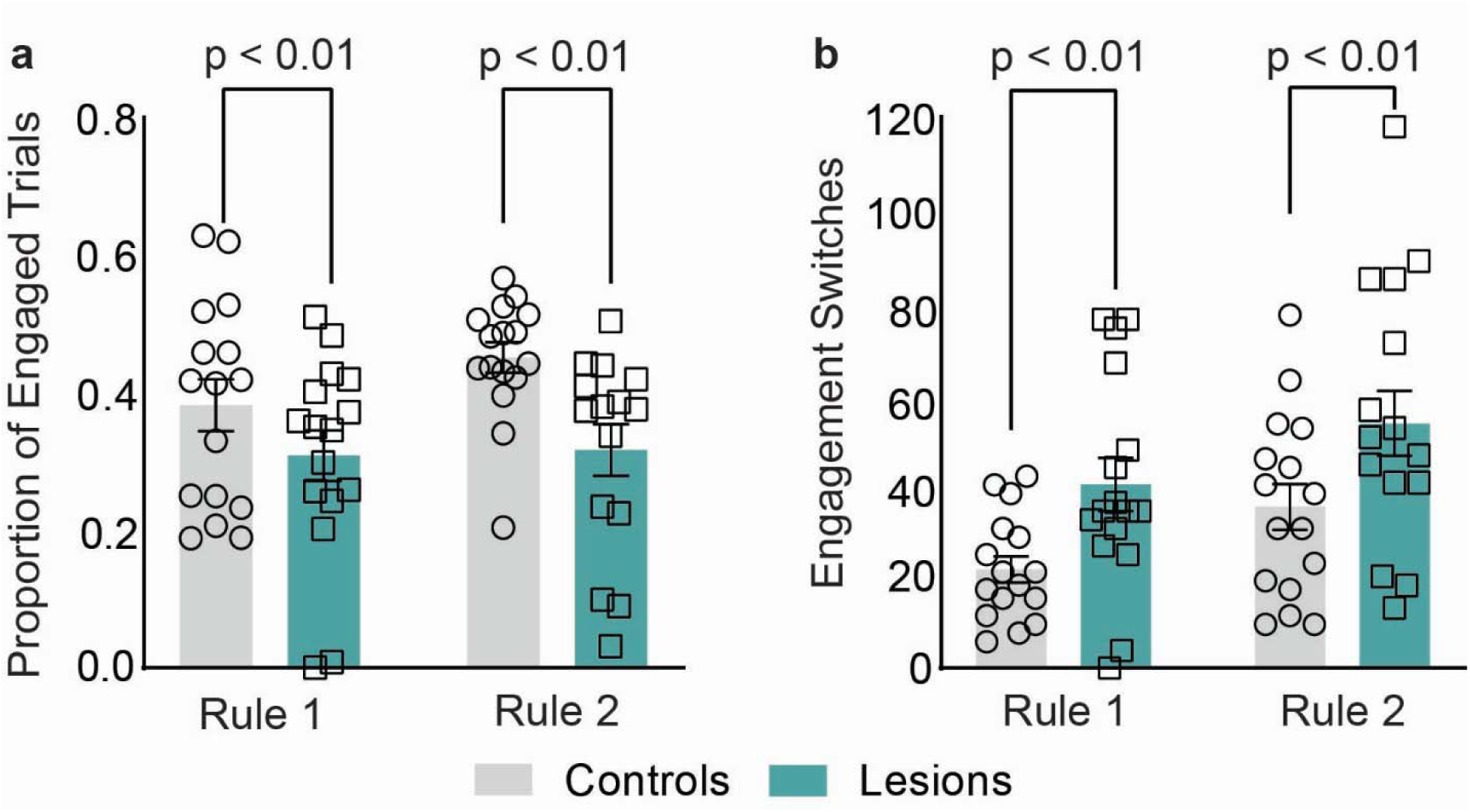
Task engagement data. a, Mean (±SEM) proportion of engaged trials in the control group and the LCN lesion group for Rules 1 and 2. b, Mean (±SEM) number of switches between engaged and non-engaged states.

These results indicate that LCN lesions caused an increase in periods of non-engagement during the categorizations tasks, which may partially explain the deficit in accuracy. Though all rats consistently alternate between engaged and non-engaged trials (Fig.6b), Lesioned rats (μ = 47.6) switched between engagement and non-engagement more often than controls (μ = 29.84), *F*(1, 59) = 10.26, *p* < 0.01. Thus, not only do LCN lesions result in proportionally less engagement in the task, rats with LCN lesions have a difficult time maintaining engagement throughout the categorization task.

### LCN lesions did not impair discrimination learning

The category learning deficit described above could have been caused by a more general deficit in visual discrimination, performance in a forced choice task, or stimulus-response mapping. We developed a control discrimination task to test for these more general deficits, which has been used in several of our previous studies [24, 27, 28]. There was no significant difference in the sessions to criterion between the lesion and control groups (Fig. 7) (*t*(14) =-1.587, *p* = 0.135), suggesting that the deficits in category learning and rule-switching were not caused by a general deficit in visual discrimination or other aspects of task performance.

**Figure 7.**
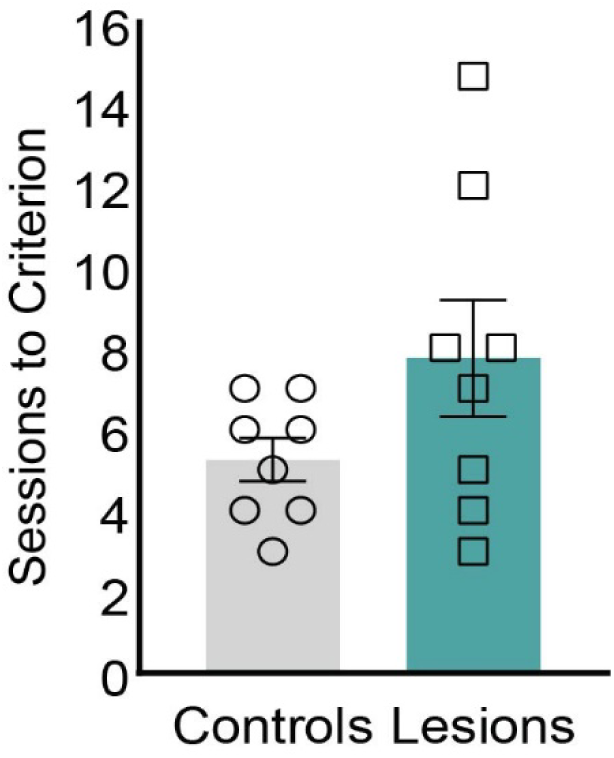
Control task data. Mean (±SEM) sessions to criterion for the control group and the LCN lesion group.

## Discussion

Our primary finding is that the cerebellum plays an important role in RB category learning. We found that LCN lesions impaired RB category learning, where rats with LCN lesions took significantly longer to reach the accuracy criterion than controls. In addition, when the category rule switched, LCN lesioned rats took longer to learn the second rule than the first rule, indicating that the cerebellum plays a role in cognitive flexibility [7–9]. Our study is the first to demonstrate a role for the cerebellum in categorization.

We assessed possible motor impairments in rats with LCN lesions to determine whether the deficits in categorization accuracy could be attributed to performance deficits or a response bias. We found that reaction times for the cue and choice phases were not affected by LCN lesions for either Rule 1 or Rule 2. The LCN lesions also did not cause a response bias. Lastly, LCN lesions did not impair simple discrimination learning, where the response requirements are identical to the RB categorization task. These findings support the hypothesis that motor and cognitive functions of the cerebellum are mediated by anatomically separate subsystems in rodents, as suggested by axonal tracing studies in primates [18, 19].

Previous work from our group indicated that the PFC focuses attention on the relevant stimulus dimension, which is crucial for RB categorization [24, 29]. In the current study, rats with LCN lesions performed similarly to rats with PFC lesions. While both groups were able to learn the categories, it took significantly longer than the controls. This finding suggests that the connection between the cerebellum and the mPFC serves an important function in categorization, and possibly other cognitive tasks. This similarity in deficits also suggests a mutual relationship, where if communication from one area is disrupted then the other area may compensate. This possible bidirectional relationship is worth investigating further to parse out how the mPFC and cerebellum work together to learn new categories. In addition, similarly to the mPFC, when the LCN was lesioned, rats did not have a deficit in the ability to discriminate between light and dark boxes in the simple discrimination task, which indicates that cerebellar communication through the LCN to mPFC is not needed for sensory discrimination, forced choice discrimination, or mapping stimuli to responses.

Control rats and humans are both able to determine and implement unidimensional categorization strategies appropriately. However, when the PFC is damaged their ability to switch from guessing or a bidimensional strategy to the appropriate unidimensional strategy is disrupted [24, 29]. In addition, they were overall less likely to use the relevant unidimensional strategy, implementing another strategy more often than controls [24, 29]. Rats with LCN lesions still switched strategies and used the relevant unidimensional strategy as much as controls. Therefore, the mPFC is important for identifying and implementing the appropriate strategy, while the cerebellum plays a different role.

Primates and rodents show transitions between task engagement and non-engagement [29, 40–42]. Our results indicate that LCN lesions caused a decrease in task engagement. In addition, we found significantly more switches between engagement states in rats with LCN lesions during both rules. These findings indicate that the cerebellum is important for maintaining the appropriate strategy. Strategy development requires selective attention whereas strategy maintenance over multiple training trials requires sustained attention. The deficits in strategy maintenance in the LCN group therefore suggests that the cerebellum plays a role in sustained attention in tasks involving numerous trials. Thus, its role contrasts with the role of the PFC which is crucial for selective attention to find the optimal strategy [24, 29].

Our use of lesions precluded the ability to determine when during training the cerebellum contributes to categorization. For instance, the cerebellum could be necessary only for initial learning and not for retention. We therefore plan to use a chemogenetic approach to elucidate when cerebellar output through the LCN is needed during category learning and retention. In addition, during this study we focused on RB categorization because the PFC is necessary for RB learning, but not information integration (II) learning. However, II learning may require LCN communication with the hippocampus and/or DMS [26–28].

## Conclusion

We found that the cerebellum plays an important role in RB category learning and category rule switching. The category learning deficit was not caused by motor deficits or a response bias. Rather, the category learning deficits with LCN lesions were related to reduced accuracy of stimulus classification, inability to maintain task engagement, and reduced cognitive flexibility. The pattern of deficits following LCN lesions suggests that the cerebellum interacts with the PFC and other forebrain systems to support multiple cognitive processes during category learning. The precise mechanisms underlying these interactions awaits physiological analyses of cerebellum-forebrain interactions.

## References

1. Buckner, R. L. (2013). The Cerebellum and Cognitive Function: 25 Years of Insight from Anatomy and Neuroimaging. Neuron, 80(3), 807–815. 10.1016/j.neuron.2013.10.044

2. Ahmadian, N., Van Baarsen, K., Van Zandvoort, M., & Robe, P. A. (2019). The Cerebellar Cognitive Affective Syndrome—A Meta-analysis. The Cerebellum, 18(5), 941–950. 10.1007/s12311-019-01060-2

3. Argyropoulos, G. P. D., Van Dun, K., Adamaszek, M., Leggio, M., Manto, M., Masciullo, M., Molinari, M., Stoodley, C. J., Van Overwalle, F., Ivry, R. B., & Schmahmann, J. D. (2020). The Cerebellar Cognitive Affective/Schmahmann Syndrome: A Task Force Paper. The Cerebellum, 19(1), 102–125. 10.1007/s12311-019-01068-8

4. Schmahmann, J. D. & Sherman, J. C. (1998) The cerebellar cognitive affective syndrome. Brain 121 (Pt 4), 561–79. 10.1093/brain/121.4.56

5. Stoodley, C.J., & Schmahmann, J. (2009). Functional topography in the human cerebellum: A meta-analysis of neuroimaging studies. NeuroImage, 44(2), 489–501. 10.1016/j.neuroimage.2008.08.039

6. Stoodley, C. J. (2012). The Cerebellum and Cognition: Evidence from Functional Imaging Studies. The Cerebellum, 11(2), 352–365. 10.1007/s12311-011-0260-7

7. Shipman, M. L., & Green, J. T. (2020). Cerebellum and cognition: Does the rodent cerebellum participate in cognitive functions? Neurobiology of Learning and Memory, 170, 106996. 10.1016/j.nlm.2019.02.006

8. Balsters, J. H., Laird, A. R., Fox, P. T., & Eickhoff, S. B. (2014). Bridging the gap between functional and anatomical features of cortico-cerebellar circuits using meta-analytic connectivity modeling: Meta-Analytic Connectivity Modeling of Cortico-Cerebellar Circuits. Human Brain Mapping, 35(7), 3152–3169. 10.1002/hbm.22392

9. Stoodley, C. J., MacMore, J. P., Makris, N., Sherman, J. C., & Schmahmann, J. D. (2016). Location of lesion determines motor vs. Cognitive consequences in patients with cerebellar stroke. NeuroImage: Clinical, 12, 765–775. 10.1016/j.nicl.2016.10.013

10. Andreasen, N. C., O’Leary, D. S., Cizadlo, T., Arndt, S., Rezai, K., Ponto, L. L., Watkins, G. L., & Hichwa, R. D. (1996). Schizophrenia and cognitive dysmetria: A positron-emission tomography study of dysfunctional prefrontal-thalamic-cerebellar circuitry. Proceedings of the National Academy of Sciences, 93(18), 9985–9990. 10.1073/pnas.93.18.9985

11. Andreasen, N.C., Paradiso, S., O’Leary, D. S. (1998). “Cognitive Dysmetria” as an Integrative Theory of Schizophrenia: A Dysfunction in Cortical-Subcortical-Cerebellar Circuitry? Schizophrenia Bulletin, 24(2), 203–218. 10.1093/oxfordjournals.schbul.a033321

12. Andreasen, N. C., & Pierson, R. (2008). The Role of the Cerebellum in Schizophrenia. Biological Psychiatry, 64(2), 81–88. 10.1016/j.biopsych.2008.01.003

13. Sugihara, I. (2018). Crus I in the Rodent Cerebellum: Its Homology to Crus I and II in the Primate Cerebellum and Its Anatomical Uniqueness Among Neighboring Lobules. The Cerebellum, 17(1), 49–55. 10.1007/s12311-017-0911-4

14. Sugihara, I., & Shinoda, Y. (2007). Molecular, Topographic, and Functional Organization of the Cerebellar Nuclei: Analysis by Three-Dimensional Mapping of the Olivonuclear Projection and Aldolase C Labeling. The Journal of Neuroscience, 27(36), 9696–9710. 10.1523/JNEUROSCI.1579-07.2007

15. Sugihara, I. (2011). Compartmentalization of the Deep Cerebellar Nuclei Based on Afferent Projections and Aldolase C Expression. The Cerebellum, 10(3), 449–463. 10.1007/s12311-010-0226-1

16. Ichinohe, N., Mori, F., & Shoumura, K. (2000). A di-synaptic projection from the lateral cerebellar nucleus to the laterodorsal part of the striatum via the central lateral nucleus of the thalamus in the rat. Brain Research, 880(1–2), 191–197. 10.1016/S0006-8993(00)02744-X

17. Pisano, T. J., Dhanerawala, Z. M., Kislin, M., Bakshinskaya, D., Engel, E. A., Hansen, E. J., Hoag, A. T., Lee, J., De Oude, N. L., Venkataraju, K. U., Verpeut, J. L., Hoebeek, F. E., Richardson, B. D., Boele, H.-J., & Wang, S. S.-H. (2021). Homologous organization of cerebellar pathways to sensory, motor, and associative forebrain. Cell Reports, 36(12), 109721. 10.1016/j.celrep.2021.109721

18. Kelly, R. M., & Strick, P. L. (2003). Cerebellar Loops with Motor Cortex and Prefrontal Cortex of a Nonhuman Primate. The Journal of Neuroscience, 23(23), 8432–8444. 10.1523/JNEUROSCI.23-23-08432.2003

19. Middleton, F. A., & Strick, P. L. (2001). Cerebellar Projections to the Prefrontal Cortex of the Primate. The Journal of Neuroscience, 21(2), 700–712. 10.1523/JNEUROSCI.21-02-00700.2001

20. Parker, K. L., Kim, Y. C., Kelley, R. M., Nessler, A. J., Chen, K.-H., Muller-Ewald, V. A., Andreasen, N. C., & Narayanan, N. S. (2017). Delta-frequency stimulation of cerebellar projections can compensate for schizophrenia-related medial frontal dysfunction. Molecular Psychiatry, 22(5), 647–655. 10.1038/mp.2017.50

21. Liu, Y., McAfee, S. S., Van Der Heijden, M. E., Dhamala, M., Sillitoe, R. V., & Heck, D. H. (2022). Causal Evidence for a Role of Cerebellar Lobulus Simplex in Prefrontal-Hippocampal Interaction in Spatial Working Memory Decision-Making. The Cerebellum, 21(5), 762–775. 10.1007/s12311-022-01383-7

22. McAfee, S. S., Liu, Y., Sillitoe, R. V., & Heck, D. H. (2019). Cerebellar Lobulus Simplex and Crus I Differentially Represent Phase and Phase Difference of Prefrontal Cortical and Hippocampal Oscillations. Cell Reports, 27(8), 2328–2334.e3. 10.1016/j.celrep.2019.04.085

23. Broschard, M. B., Kim, J., Love, B. C., & Freeman, J. H. (2021). Category learning in rodents using touchscreen-based tasks. Genes, Brain and Behavior, 20(1), e12665. 10.1111/gbb.12665

24. Broschard, M. B., Kim, J., Love, B. C., Wasserman, E. A., & Freeman, J. H. (2021). Prelimbic cortex maintains attention to category-relevant information and flexibly updates category representations. Neurobiology of Learning and Memory, 185, 107524. 10.1016/j.nlm.2021.107524

25. Broschard, M. B., Kim, J., Love, B. C., Wasserman, E. A., & Freeman, J. H. (2019). Selective attention in rat visual category learning. Learning & Memory, 26(3), 84–92. 10.1101/lm.048942.118

26. Broschard, M. B., Kim, J., Halverson, H. E., Farley, S. J., & Freeman, J. H. (2025) Interaction between the medial prefrontal cortex, dorsomedial striatum, and dorsal hippocampus that support rat category learning. bioRxiv. 10.1101/2025.04.21.649785

27. Broschard, M. B., Kim, J., Love, B. C., & Freeman, J. H. (2023). Dorsomedial striatum, but not dorsolateral striatum, is necessary for rat category learning. Neurobiology of Learning and Memory, 199, 107732. 10.1016/j.nlm.2023.107732

28. Broschard, M. B., Kim, J., Love, B. C., Halverson, H. E., & Freeman, J. H. (2024). Disrupting dorsal hippocampus impairs category learning in rats. Neurobiology of Learning and Memory, 212, 107941. 10.1016/j.nlm.2024.107941

29. Broschard, M.B., Turner, B., Tranel, D., & Freeman, J.H. (2024). Dissociable roles of the dorsolateral and ventromedial prefrontal cortex in human categorization. Journal of Neuroscience, 44, doi: 0.1523/JNEUROSCI.2343-23.2024.

30. Logan, G. D., Zbrodoff, N. J., & Williamson, J. (1984). Strategies in the color-word Stroop task. Bulletin of the Psychonomic Society, 22(2), 135–138. 10.3758/BF03333784

31. Ell S. W. & Ivry, R. B. (2008). Cerebellar pathology does not impair performance on identification or categorization tasks. Journal of the Internation Neuropsychological Society, 14, 760–770. doi:10.10170S1355617708081058

32. Maddox, W. T., Aparicio, P., Marchant, N. L., Ivry, R. B. (2005). Rule-based category learning is impaired in patients with Parkinson’s disease but not in patients with cerebellar disorders. Journal of Cognitive Neuroscience, 17(5), 707–723. doi:10.1162/0898929053747630

33. Crijns, E., & Op de Beeck, H. (2019). The Visual Acuity of Rats in Touchscreen Setups. Vision, 4(1), 4. 10.3390/vision4010004

34. Jaeger, T. F. (2008). Categorical data analysis: Away from ANOVAs (transformation or not) and towards logit mixed models. Journal of Memory and Language, 59(4), 434–446. 10.1016/j.jml.2007.11.007

35. Hélie, S., Turner, B. O., Crossley, M. J., Ell, S. W., & Ashby G. F. (2017). Trial-by-trial identification of categorization strategy using iterative decision-bound modeling. Behav Res 49, 1146–1162. 10.3758/s13428-016-0774-5

36. O’Donoghue, E. M., Broschard, M. B., Freeman, J. H., & Wasserman, E. A. (2022). The Lords of the Rings: People and pigeons take different paths mastering the concentric-rings categorization task. Cognition, 218, 104920. 10.1016/j.cognition.2021.104920

37. Neath, A. A., & Cavanaugh, J. E. (2011). The bayesian information criterion: Background, derivation, and applications. WIREs Computational Statistics, 4(2), 199–203. 10.1002/wics.199

38. Vuong, Q. H. (1989). Likelihood ratio tests for model selection and non-nested hypotheses. Econometrica, 57(2), 307. 10.2307/1912557

39. Paxinos, G. and Watson, C. (2007) The Rat Brain in Stereotaxic Coordinates. 6th Edition, Academic Press, San Diego.

40. Ashwood, Z. C., Roy, N. A., Stone, I. R., Urai, A. E., Churchland, A. K., Pouget, A., & Pillow, J. W. (2020). Mice alternate between discrete strategies during perceptual decision-making. Nature Neuroscience, 25, 201–212. 10.1101/2020.10.19.346353

41. Pettit, N.L., Yuan, X.C., & Harvey, C.D. (2022). Hippocampal place codes are gated by behavioral engagement. Nature Neuroscience, 25, 561–566. 10.1038/s41593-022-01050-4

42. Grohn, J., et al., (2024). General mechanisms of task engagement in the primate frontal cortex. bioRxiv. 10.1101/2024.01.16.575830

